# Interaction between VPS13A and the XK scramblase is required to prevent VPS13A disease in humans

**DOI:** 10.1101/2022.05.09.491223

**Authors:** Jae-Sook Park, Yiying Hu, Nancy. M. Hollingsworth, Gabriel Miltenberger-Miltenyi, Aaron M. Neiman

## Abstract

VPS13 family proteins form conduits between the membranes of different organelles through which lipids are transferred. In humans there are four VPS13 paralogs each of which is required to prevent a different inherited disorder. VPS13 proteins contain multiple conserved domains. The VAB domain binds to adaptor proteins to recruit VPS13 to specific membrane contact sites. This work demonstrates the importance of a different domain in VPS13A in preventing VPS13A disease (chorea-acanthocytosis). The Pleckstrin Homology (PH) domain at the C-terminus of VPS13A is required to form a complex with the XK scramblase and for proper localization of VPS13A within the cell. Mutations in the interaction surface between VPS13A and XK predicted by Alphafold modeling disrupt complex formation and colocalization of the two proteins. Mutant *VPS13A* alleles found in patients with VPS13A disease truncate the PH domain. The phenotypic similarities between VPS13A disease and McLeod syndrome caused by mutations in *XK* argue that loss of VPS13A-XK complex is the basis of both diseases.

**Summary Statement:** VPS13A disease and McLeod syndrome are related disorders caused by mutation of the *VPS13A* and *XK* genes, respectively. A pathologic *VPS13A* mutation disrupts binding of the VPS13A and XK proteins, suggesting a common basis of both diseases.

## Introduction

In humans there are four VPS13 paralogs, *VPS13A* through *D*. Mutations in each *VPS13* gene are associated with different inherited disorders (Gauthier et al., 2018; Kolehmainen et al., 2003; Lesage et al., 2016; Rampoldi et al., 2001; Seong et al., 2018; Ueno et al., 2001). For example, the adult-onset neurodegenerative disease chorea-acanthocytosis (also known as VPS13A disease) is due to loss of *VPS13A* function (Rampoldi et al., 2001; Ueno et al., 2001; Walker and Danek, 2021). VPS13 family members belong to an important class of lipid transfer proteins that localize to membrane contact sites created where membranes of different organelles come into close apposition (Dziurdzik and Conibear, 2021; Leonzino et al., 2021; McEwan and Ryan, 2021; Melia and Reinisch, 2022). For example, VPS13A is present at both endoplasmic reticulum (ER)-mitochondrial junctions and ER-lipid droplet junctions (Kumar et al., 2018; Munoz-Braceras et al., 2019; Yeshaw et al., 2019). The rod-shaped VPS13 protein spans the gap between two organelles and a hydrophobic groove running through the interior of the protein allows lipids to transit in bulk between the two membranes (Leonzino et al., 2021; Li et al., 2020).

In yeast, there is a single Vps13 protein that localizes at different membrane contact sites depending on the growth and differentiation state of the cell (Lang et al., 2015; Park et al., 2016). The distribution of Vps13 to these different sites is regulated by its interaction with organelle-specific adaptor proteins that compete for binding to a region of Vps13 termed the Vps13 adaptor binding (VAB) domain (Figure 1A)(Bean et al., 2018). Loss of a particular adaptor protein releases Vps13 from specific organelles resulting in a subset of phenotypes observed in *vps13* null mutants (Lang et al., 2015; Park et al., 2013).

**Figure 1.**
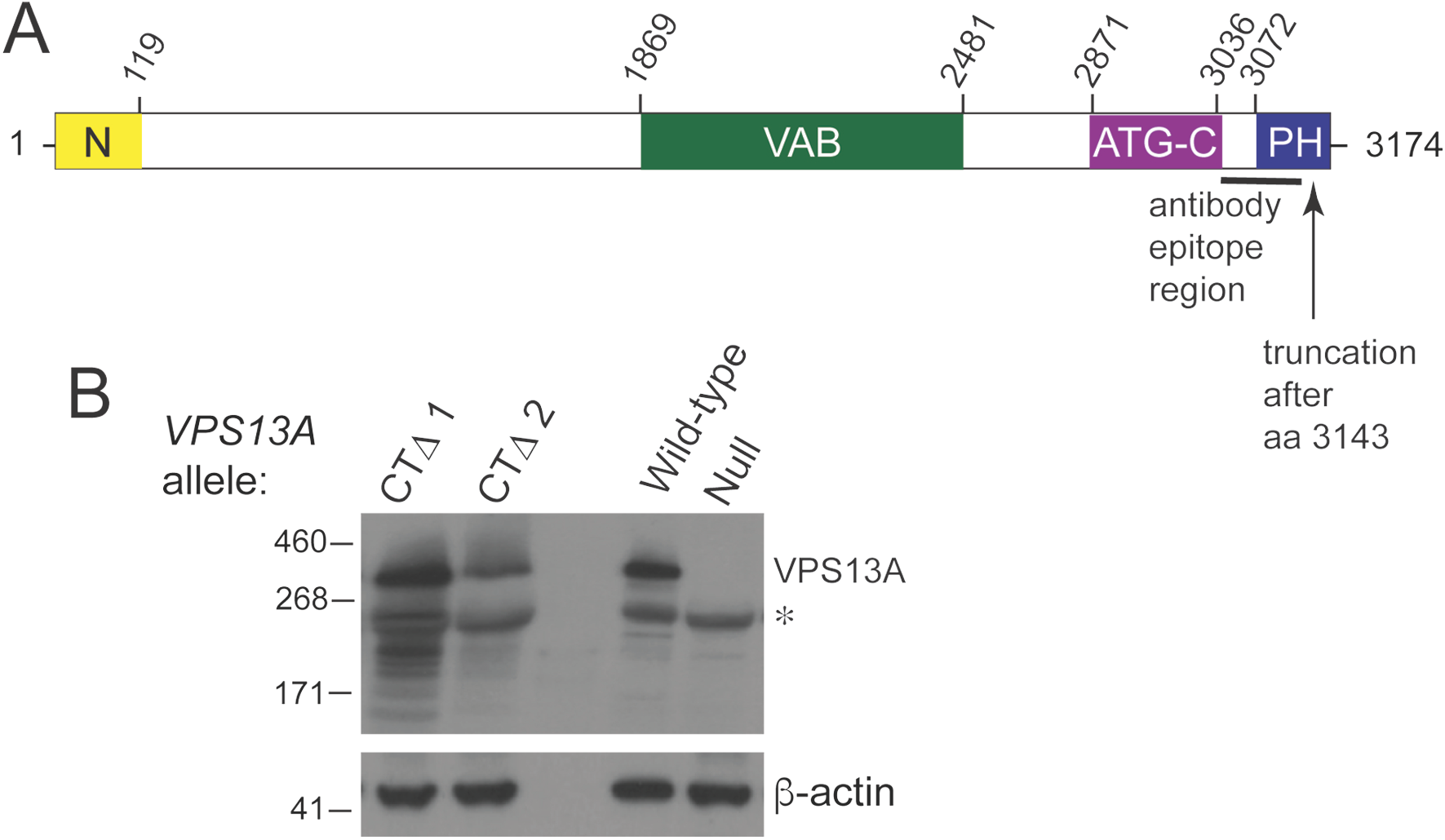
Truncation of the VPS13A C-terminus creates a stable protein and VPS13A disease. (A) Schematic of the VPS13A protein. The domain arrangement of VPS13A. Yellow indicates the N-Chorein domain, green represents the Vps13 adaptor binding (VAB) domain, purple is the ATG2 C-terminal domain, and blue denotes the PH-like domain (Rzepnikowska et al., 2017). Numbers above indicate the amino acid positions of the domains. The black line indicates the position of the protein fragment used to produce the VPS13A antibody used in (B), while the arrow indicates the end of the CTΔ truncation. (B) Immunblot analysis of VPS13A proteins from human blood samples. Samples from patients with VPS13A disease were probed on immunoblots with α-VPS13A antibodies that recognize an epitope in the C-terminal region of the protein. CTΔ #1 and CTΔ #2 indicate samples from Patients 1 and 2, respectively. WT is a control sample from a healthy individual, and “null” is a sample from a patient diagnosed with VPS13A disease that fails to exhibit the protein {Niemela, 2020 #1226). β-actin was used as a loading control. The asterisk indicates a prominent cross-reacting species. β-actin was used as a loading control.

The yeast paradigm likely holds true for the VPS13 paralogs in human cells. Some patients with VPS13A disease have missense mutations in the VPS13A VAB domain indicating that VPS13A binding to adaptor proteins is important for its function (Dobson-Stone et al., 2002). One candidate for a VPS13 adaptor protein is XK. XK is a “scramblase” that is able to flip lipids between leaflets of a lipid bilayer (Adlakha et al., 2022; Ryoden et al., 2022). People with mutations in the *XK* gene exhibit a neurodegenerative disorder known as McLeod syndrome that shares phenotypic similarities with VPS13A disease (Roulis et al., 2018). The XK protein co-localizes with VPS13A in cells by formation of a VPS13A-XK complex (Park and Neiman, 2020; Urata et al., 2019). However a VAB domain mutant did not disrupt formation of the VPS13A-XK complex *in vivo* (Park and Neiman, 2020). Since VAB domain mutants may disrupt binding to multiple different adaptor proteins, the question of whether VPS13A and XK function together remained an open question. This work shows that XK forms a complex with VPS13A, not through binding to the VAB domain, but instead to a PH domain localized at the C-terminal end of VPS13A. Furthermore, we demonstrate that two unrelated patients with VPS13A disease carry an allele of *VPS13A* that encodes a stable protein truncating this domain and disrupting interaction with XK. These results provide *in vivo* evidence that VPS13A disease and McLeod syndrome share as a common basis the loss of functional VPS13A-XK complexes.

## Results

### A small truncation of the VPS13A C-terminus results in VPS13A disease

Two unrelated patients with VPS13A disease both carry the *vps13A-c9431_9432delAG* allele (hereafter *VPS13A-CT*Δ for C-Terminal Deletion) (Figure 1A) (Dobson-Stone et al., 2002; Richard et al., 2019). This allele contains a frameshift mutation that removes the last 31 amino acids (aa) from the end of the 3174aa VPS13A protein while adding six additional amino acids to the C-terminus. The second alleles in Patient 1 and Patient 2 are *vps13A-c.6725delA* and *vps13A-c.4956+1G>A*, respectively (Dobson-Stone et al., 2002; Richard et al., 2019). These alleles contain a nonsense and a splice site mutation, respectively that could produce only severely truncated protein (Dobson-Stone et al., 2002). To determine whether the VPS13A-CTΔ protein is stably expressed, erythrocyte membrane samples from Patients 1 and 2 were analyzed by immunoblots using α-VPS13A antibodies that recognize the C terminal region of VPS13A (Figure 1A). The VPS13A positive control came from a healthy individual, while the negative control used membrane samples from a patient with VPS13A disease in which the protein was known to be absent {Niemela, 2020 #1226). Both patients carrying the *VPS13A-CT*Δ allele displayed VPS13A protein with similar mobility to WT, as expected given that the 3 kDa lost in the truncation comprises a small fraction of the total 349 kDa WT protein (Figure 1B). The extreme C-terminal region of VPS13A is therefore critical for a function in humans that prevents development of VPS13A disease.

### The VPS13A C-terminus is important for cellular localization and VPS13A-XK complex formation

The *vps13A-ΔCT* allele affects the C-terminus is two ways: (1) removal of the last 31 amino acids and (2) addition of six amino acids to the end of the truncated protein. To see whether it is the truncation that affects VPS13A function, a stop codon was introduced immediately after codon 3143 in an allele of *VPS13A* that carries an internal fusion of the mCherry fluorescent protein gene (designated *VPS13A^mCherry*) {Kumar, 2018 #1030}. This mutant allele, as well as a WT control, was transfected into the human embryonic kidney cell line, HEK293T, and the VPS13A proteins visualized by fluorescence microscopy.

Previous work showed that VPS13A localizes predominantly to short patches throughout the ER, representing ER-mitochondrial contacts, and on lipid droplets (Kumar et al., 2018; Munoz-Braceras et al., 2019; Yeshaw et al., 2019). Consistent with these results, VPS13A^mCherry localization to lipid droplets was quite prominent (100% of lipid droplets displaying VPS13A^mCherry fluorescence, >100 cells scored; Figure 2A). In contrast the VPS13A^1-3143^^mCherry protein exhibited only a diffuse cytosolic fluorescence and co-localization with lipid droplet marker was greatly reduced (1.8% of lipid droplets displaying VPS13A^mCherry fluorescence, >100 cells scored; Figure 2A).

**Figure 2.**
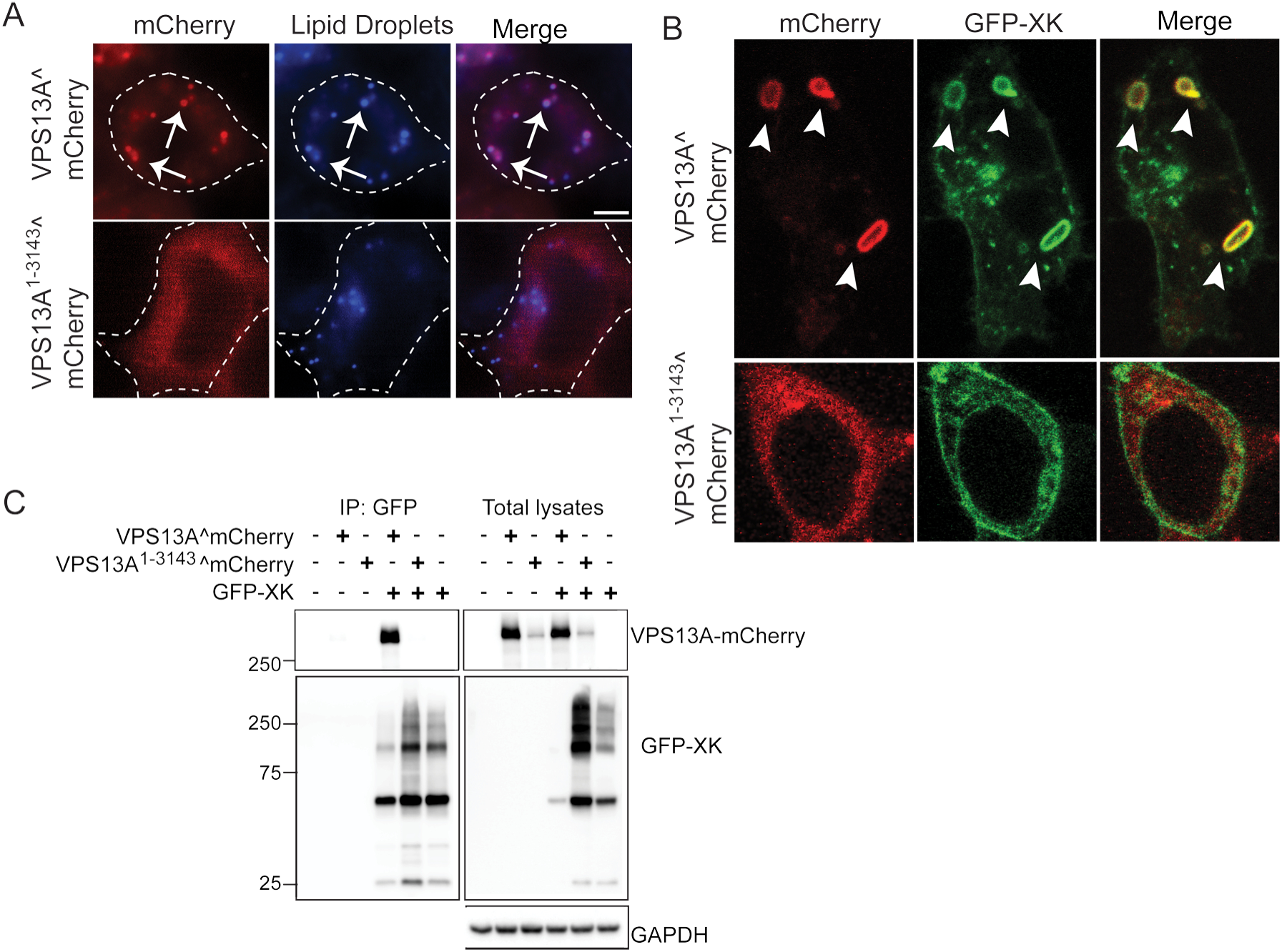
The VPS13A C-terminus is required for co-localization with XK in cells and for VPS13A-XK complex formation. **(A)** Cellular localization. HEK293T cells were transfected with plasmids expressing either *VPS13A^mCherry* (p*VPS13A*^mCherry) or *VPS13A^1-3143^^mCherry* (pJS160). Protein localization was examined by wide field fluorescence microscopy. Lipid droplets were visualized using the blue fluorescent marker monodansylpentane (MDH){Currie, 2014 #972}. Arrows highlight a subset of examples of co-localization between VPS13A^mCherry and lipid droplets. Scale bar = 5 μm. (B) HEK293T cells were co-transfected with plasmids expressing either *VPS13^mCherry* or *VPS13A^1-3143^^mCherry* together with *GFP-XK*. Arrowheads indicate co-localization of VPS13^mCherry with GFP-XK. Scale bar = 5 μm. (C) Co-immunoprecipitation. HEK293T cells were transfected with either *VPS13A^mCherry* or *VPS13A^1-3143^^mCherry* with or without *GFP-XK*. Twenty-four hours after transfection, cells were lysed and GFP nanobodies coupled to agarose beads were used to isolate the GFP-XK. Bound proteins were probed on immunoblots with α-GFP and α-mCherry antibodies to detect XK and VPS13A, respectively. Right panels shows total lysates prior to incubation with the GFP nanobodies. GAPDH was used as a loading control.

Colocalization of VPS13A^mCherry was altered when *GPF-XK* was simultaneously over-expressed, moving from lipid droplets to patches in the ER where it co-localized with the XK protein (100% colocalization, >25 cells examined in each of three independent experiments; Figure 2B) (Park and Neiman, 2020). However, when *VPS13A^1-3143^^mCherry* was co-transfected with *GFP-XK*, no co-localization with XK was observed (0% co-localization, >25 cells examined in each of three independent experiments; Figure 2B). These results indicate that the last 31 amino acids of VPS13A are required for VPS13A-XK co-localization with the cell.

To see if the failure of the VPS13A and XK proteins to colocalize was due to a defect in VPS13A-XK complex formation, co-immunoprecipitation (co-IP) experiments were performed. *GFP-XK* was co-transfected into HEK293T cells with either *VPS13A^mCherry* or *VPS13A^1-3143^^mCherry* and the GFP-XK protein was precipitated from cell lysates using anti-GFP nanobodies. These precipitates were then probed with α-mCherry antibodies to detect VPS13A. As a specificity control, cells transfected with only the *VPS13A^mCherry* constructs were also tested. VPS13A^mCherry protein was present in the co-IP with GFP-XK but not when GFP-XK was absent (Figure 2C). No co-IP was observed for GFP-XK and VPS13A^1-3143^^mCherry (Figure 2C). While the steady state VPS13A^1-3143^^mCherry protein level was reduced in total lysates, this is likely not the cause of the failure to co-IP with GFP-XK as significantly more GFP-XK was present in the precipitate (Figure 2C). VPS13A-XK complex formation, therefore, occurs via interaction with the C-terminal 31 aa of VPS13A and is required for cellular co-localization.

### The C-terminal PH domain of VPS13A is sufficient for XK binding

The final 31 aa of VPS13A are part of a Pleckstrin Homology (PH) domain, a common structural motif known to be involved in both protein-protein and protein-lipid interactions (Scheffzek and Welti, 2012). To determine whether the intact PH domain is required for XK interaction or just the extreme C-terminus, GFP fusions to different parts of the VPS13A C-terminus were constructed. *GFP-VPS13A^2751-3174^* includes the ATG-C and PH domains (Figure 1A), *GFP-VPS13A^3927-3174^*contains only the intact PH domain, *GFP-VPS13A^3027-3143^* has the PH domain with the 31 aa truncation and *GFP-VPS13A^3144-3174^* has only the last 31 aa. These alleles were transfected into HEK293T cells with and without *XK* and steady state VPS13A protein levels examined by immunoblots using α-GFP antibodies. The three larger fusions exhibited similar protein levels regardless of the presence or absence of *XK* (Figure 3A). The smallest fusion could not be resolved from GFP alone, and therefore proteolytic cleavage of the 31aa from the GFP cannot be ruled out.

**Figure 3.**
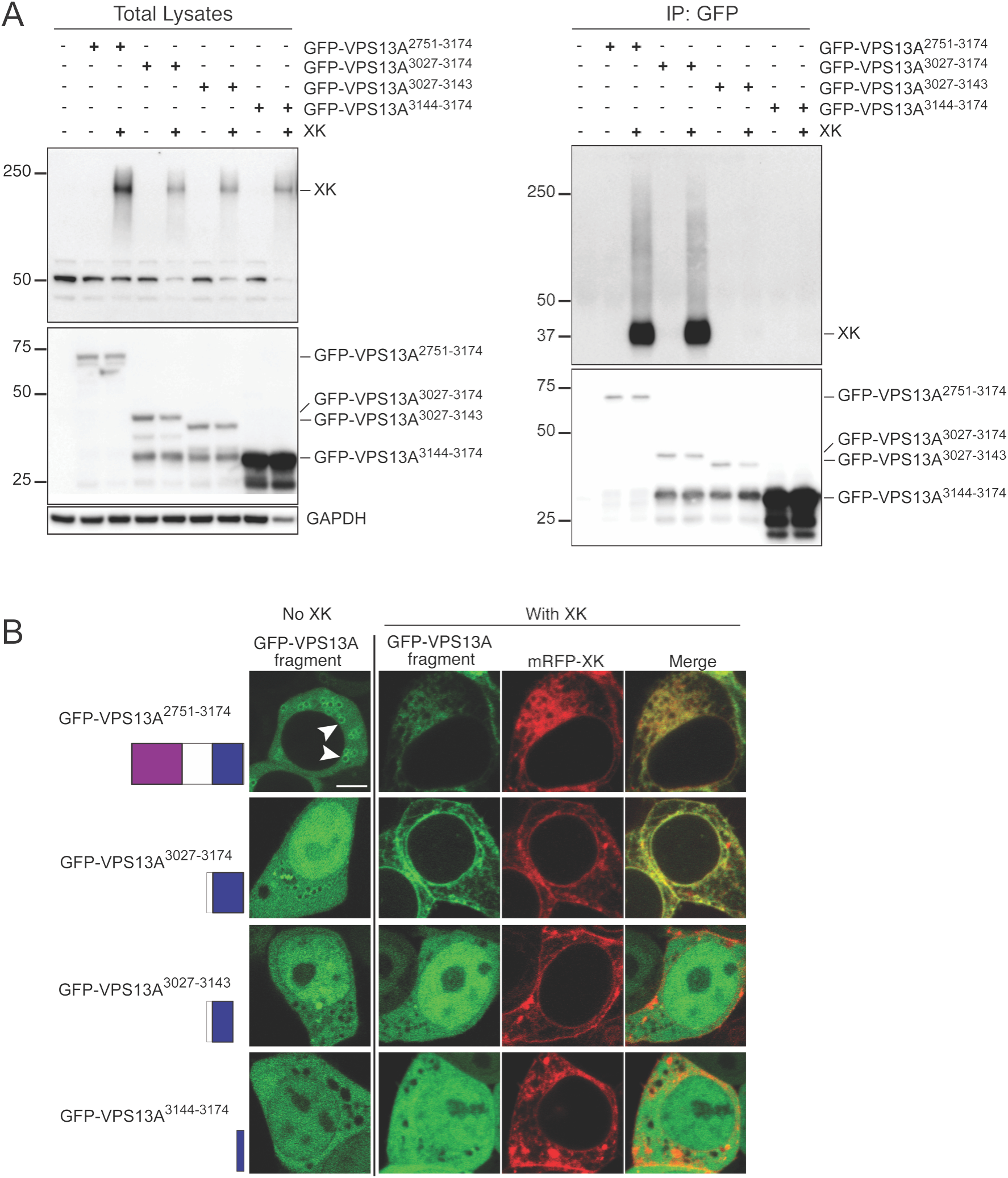
The VPS13A PH domain is sufficient for VPS13A-XK complex formation. (A) Coimmunopreciptition of XK with different fragments of VPS13A fused to GFP. HEK293T cells were transfected with plasmids expressing *GFP-VPS13A^2751-3174^* (pJS164) that contains the ATG-C (purple box) and PH (blue box) domains, *GFP-VPS13A^3027-3174^* (pJS166) that contains only the PH domain, *GFP-VPS13A^3027-3143^*(pJS171) that contains the 31 aa acid truncation of the PH domain, or *GFP-VPS13A^3144-3174^* (pJS167) that contains on the last 31 aa of VPS13A, with or without co-transfections with a vector expressing an untagged *XK* gene (pJS169). Twenty-four hours after transfection, cells were lysed and total lysates were probed on immunoblots with α-XK antibodies α-GFP antibodies to detect different GFP-VPS13A fusions. α -GAPDH antibodies were used as loading control. Nanobodies coupled to agarose beads were used to isolate the GFP-VPS13A fusion proteins and the precipitates were then probed as for the total lysates. (B) Localization. HEK293T cells were transfected with the same combinations of vectors as in (A) and then protein localization was examined by confocal fluorescence microscopy. Arrowheads in the upper left panel indicate examples of enrichment of GFP fluorescence on the lipid droplet surface. When expressed alone, only VPS13A^2751-3174^ displayed this concentration on the lipid droplet surface (69% of cells vs 0% for all other constructs >100 cells scored in two independent experiments). Residue numbers and representation of the domains is shown at left. Scale bar = 5μm.

Fluorescence microscopy was used to examine the localization patterns of the different fusions in presence or absence of transfected *mRFP-XK*. In the absence of exogenous *mRFP-XK*, GFP-VPS13A^2751-3174^ localized diffusely in the cell with some concentration on the surface of lipid droplets (arrowheads in Figure 3B), as described previously(Kumar et al., 2018). This concentration was lost when *mRFP-XK* was overexpressed, with GFP-VPS13A^2751-3174^ instead mirroring the distribution of mRFP-XK throughout the endomembrane system (Figure 3B). This is similar to the behavior of full-length VPS13A^mCherry in that overexpression of *XK* competes VPS13A away from lipid droplets to the ER(Park and Neiman, 2020). Thus, the C-terminal 424 aa of VPS13A contain the region essential for XK association but other regions are necessary to limit that colocalization to the ER.

The GFP-VPS13^3027-3174^ C-terminal fusion containing only the PH domain localized diffusely throughout both the cytoplasm and nucleoplasm without any clear concentration on lipid droplets (Figure 3B). Thus, the ATG-C domain is necessary for lipid droplet localization, consistent with the results of Kumar *et al*. (2018). When co-expressed with *mRFP-XK*, the distribution of GFP-VPS13^3027-3174^ within the cell changed to mirror XK, similar to what was observed for GFP-VPS13A^2751-3174^ (Figure 3B). Therefore, the PH domain of VPS13A is sufficient for association with XK. Removal of the last 31 aa from the PH domain abolished co-localization with XK, as did the fusion with just the 31 aa alone (Figure 3B). These results show that the C-terminal 31 aa of VPS13A are necessary but not sufficient for co-localization with XK.

To determine if these co-localization defects reflect changes in the ability of the VPS13A fragments to bind to XK, the same constructs were transfected into HEK293T cells with or without a vector expressing untagged *XK*. The GFP fusions were precipitated and the co-precipitation of XK assessed using anti-XK antibodies (Figure 3C. Note that endogenous XK protein levels are too low to be detected under these conditions either in total lysates or the GFP-VPS13A precipitates (Figure 3A, 3C, lanes 1 and 2). Consistent with the localization results, overexpressed XK protein co-precipitated with both GFP-VPS13A^2751-3174^ and GFP-VPS13^3027-3174^ but with neither of the two shorter fragments. For reasons that are unclear, the mobility of the XK protein in gels was reproducibly faster in the precipitated samples than in the total lysates. The observed ∼37 kDa molecular weight of this faster migrating XK species is closer to that expected for the predicted molecular weight of XK (51 kDa) than the ∼200 kDa protein detected in the total lysates. One possible explanation for the slow mobility of XK in total lysates is that the protein was not completely solubilized in these samples. Together with the localization data, these results demonstrate that the PH domain at the C-terminus of VPS13A is the region responsible for binding to the XK protein.

### Mutations in the second intracellular loop of XK disrupt binding to the VPS13A PH domain

XK is an integral membrane protein with 10 predicted hydrophobic regions. An Alphafold structural prediction of XK suggests that two of these hydrophobic stretches are intramembrane hairpins rather than transmembrane domains as was previously proposed (Figure 4A) ((Jumper et al., 2021; Varadi et al., 2022). A recent cryo-EM structure of the related protein XKR8 supports this topological arrangement for XK(Ryoden et al., 2022). The consequence of this change is that three loops previously predicted to be extracellular loops are actually cytoplasmic. XKR2 encodes the most closely related paralog of XK in human cells and co-precipitated with VPS13A in a proteomic study (Huttlin et al., 2015; Suzuki et al., 2014). On the assumption that the the VPS13A binding site is conserved between XKR2 and XK, the cytoplasmic loops of these two proteins were aligned, revealing an identical six amino acid sequence in the second intracellular loop of both proteins (Figure 4B). Based on this alignment, an allele of *XK* was constructed in which these six conserved residues (EEPYVS at aa 97-102 of XK) were mutated to alanine ((*XK^6A^*).

**Figure 4.**
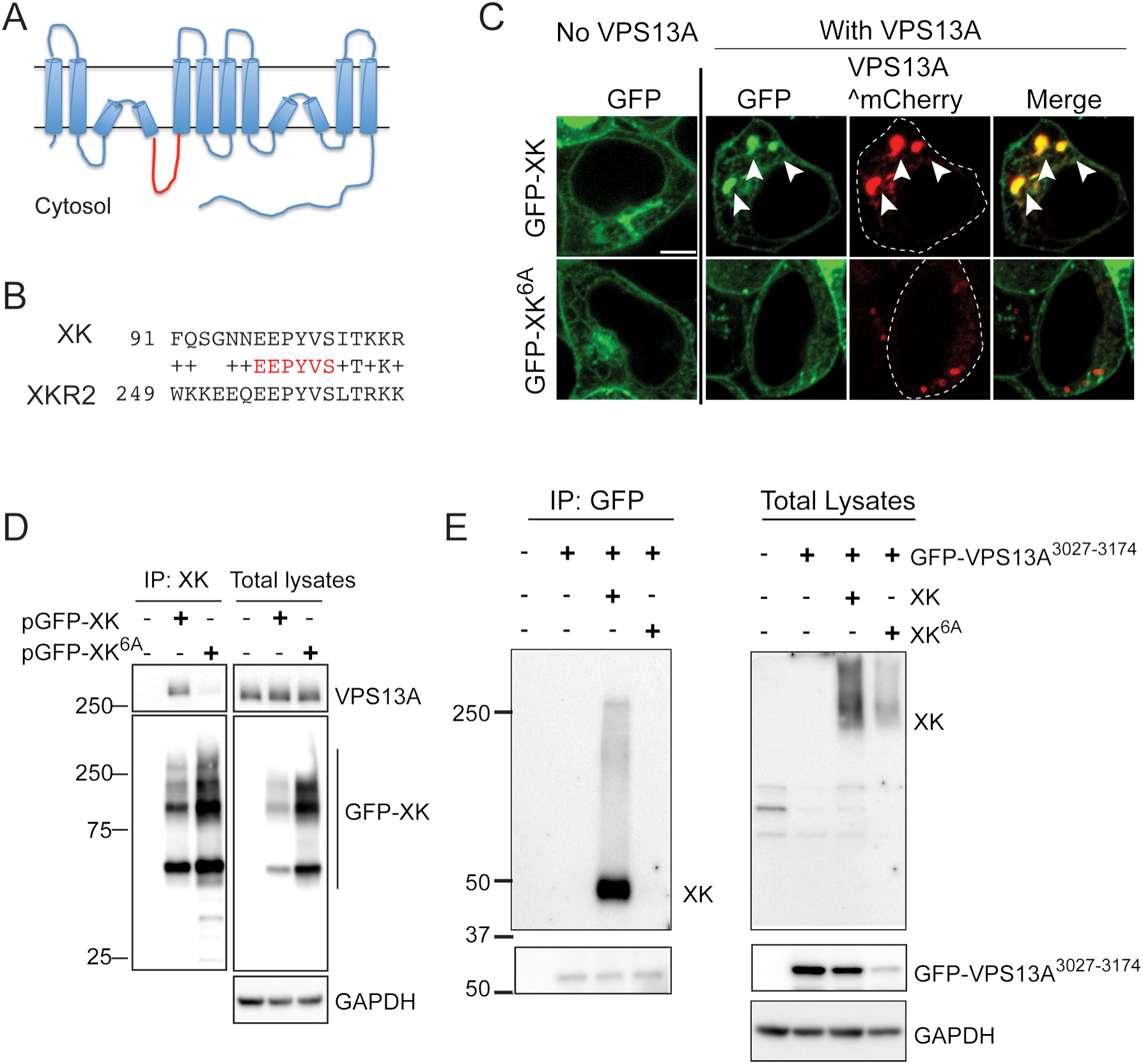
Amino acids in the second intracellular loop of XK are required for interaction with VPS13A. (A) Predicted topology of the XK protein based on an Alphafold structural prediction (Jumper et al., 2021; Varadi et al., 2022). The second intracellular loop is highlighted in red. (B) Alignment of sequences from the second intracellular loop of XK and XKR2. The invariant residues highlighted in red were mutated to alanines in the *XK-6A* allele. (C) Localization. HEK293T cells were transfected with constructs expressing either *GFP-XK* or *GFP-XK^6A^* with or without *VPS13A*^mCherry. Arrowheads highlight co-localization of GFP-XK and VPS13A^mCherry. Scale bar = 5μm. (D) Co-immunoprecipitation of endogenous VPS13A with GFP-XK or GFP-XK^6A^. HEK293T cells were transfected with constructs expressing either GFP-XK (pcDNA3.1(+)-N-eGFP-*XK*) or GFP-XK^6A^ (pJS165). GFP-XK was precipitated with GFP nanobodies as in Figure 2C. Precipitates and total lysates were probed on immunoblots using α-VPS13A and α-GFP antibodies (left panels). GAPDH was used as a loading control. (E) Co-immunoprecipitation of overexpressed untagged *XK* or *XK^6A^* with the VPS13A PH domain. HEK293T cells were transfected with a construct expressing the GFP-VPS13A^3027-3174^ fusion alone (pJS166) with untagged XK (pJS169) or XK^6A^ (pJS173) and processed as described in Figure 3A.

HEK293T cells expressing *GFP-XK^6A^* in the presence of endogenous *VPS13A* displayed similar levels of fluorescence and subcellular distribution as the WT GFP-XK protein (Figure 4C). When *VPS13A^mCherry* was co-expressed with *GFP-XK*, 100% of the cells exhibited co-localization of the two proteins (Figure 4C) (n > 140 cells over three independent experiments). This number was reduced to just 9% when *GFP-XK^6A^* was combined with *VPS13A^mCherry* (n > 350 cells scored over five independent experiments).

The conserved residues in the second intracellular loop not only promote VPS13A^mCherry-XK co-localization, they are also required for VPS13A-XK complex formation. Endogenous VPS13A co-precipitated with GFP-XK, but not GFP-XK^6A^, even though more of the mutant protein was present in the precipitate (Figure 4D) (Figure 4D). Furthermore, this region of XK specifically interacts with the VPS13A PH domain as untagged XK^6A^ failed to co-precipitate with the GFP-VPS13^3027-3174^ fragment (Figure 4E).

### Alphafold prediction of the VPS13A-XK binding interface

To define more precisely the binding interface between VPS13A and XK, the multimer mode of Alphafold2 was used to predict the structure of the PH domain of VPS13A in complex with the second intracellular loop of XK (Figure 5A) (Evans et al., 2022; Jumper et al., 2021). The XK sequence is largely unstructured except for two short β strands that align with three β strands of the PH domain in VPS13A to create a short β-sheet. Thus, the predicted interface between the two proteins is the contact between one β strand from XK and one from VPS13A. Our results above are consistent with this model as the 31aa truncation of VPS13A that disrupts the XK interaction removes this critical β strand in VPS13A. Similarly, the mutations present in the *XK^6A^*allele mutate three residues in the β strand of XK that aligns with VPS13A. To test the prediction more directly, the *I3148P* mutation was introduced into *GFP-VPS13^3027-3174^*. This mutation is predicted to introduce a twist in the VPS13A β strand that could disrupt interaction with XK. The wild-type and I3148P versions of *GFP-VPS13^3127-3174^* were both expressed in HEK293T cells and examined for colocalization with mRFP-XK. While both VPS13A proteins localized throughout the cell when expressed alone, the subcellular localization of the GFP-VPS13A^3217-3174^-I3148P did not change in the presence of mRFP-XK, unlike GFP-VPS13A^3217-3174^ (Figure 5B). Similarly, introduction of the I3148P mutation in the GFP-VPS13^3027-3174^ protein abolished the co-IP with XK without affecting the stability of GFP-VPS13^3027-3174^ (Figure 5C). Thus, this point mutation was sufficient to disrupt the interaction between the VPS13A and XK, providing additional support for the Alphafold2 model of the binding interface.

**Figure 5.**
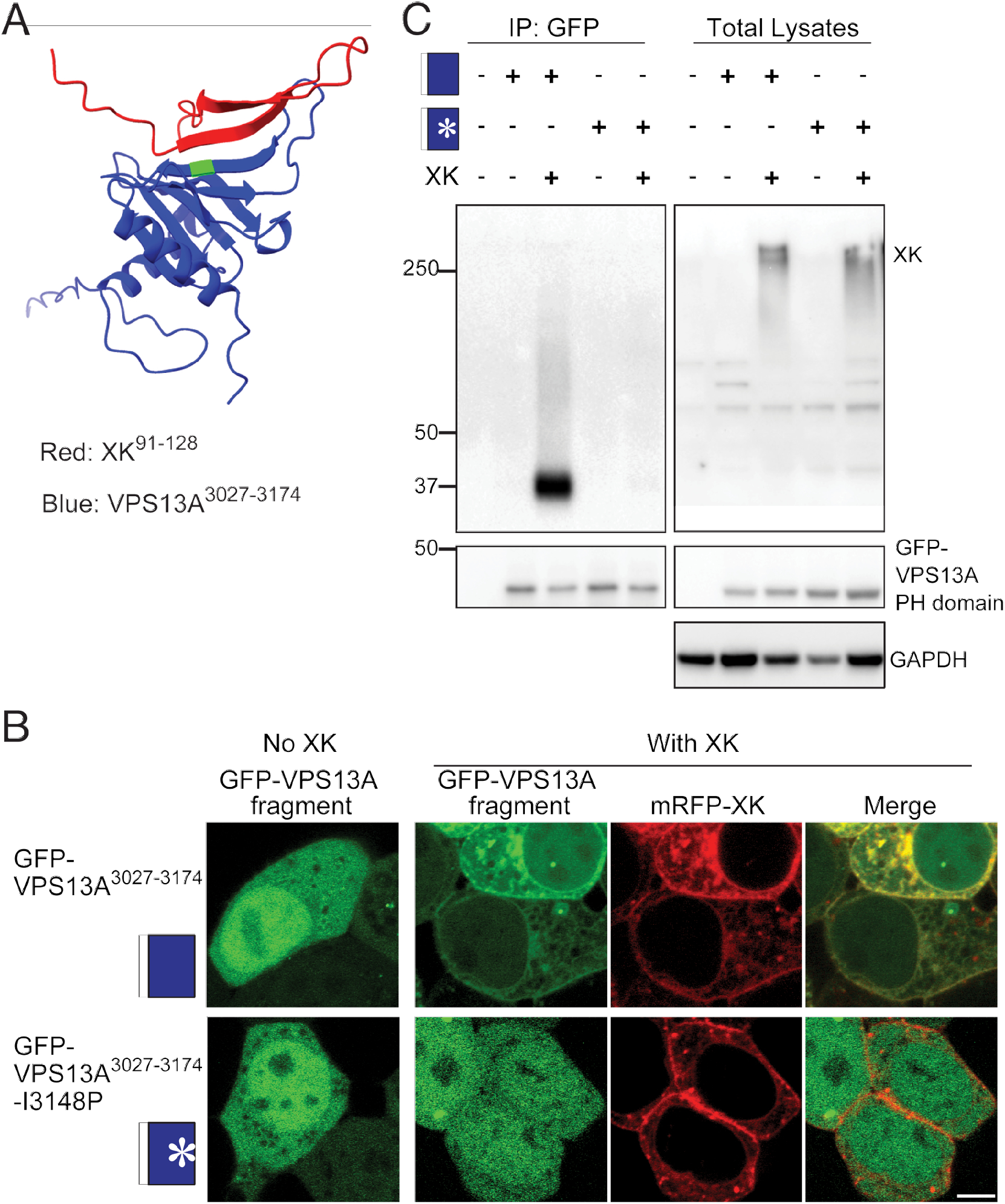
Identification of an XK-VPS13A binding interface. (A) Alphafold multimer assembly of the VPS13A PH domain (blue) with the second intracellular loop of XK (red) showing antiparallel alignment of a predicted ß-strand in XK (aa 100-106) with one in VPS13A (aa 3144-3150). Green shows the position of I3148 of VPS13A. (B) Cellular localization. The localization of GFP-VPS13A^3027-3174^ or GFP-VPS13A^3027-3174^-I3148P (pJS172) was analyzed in HEK293T cells with and without co-transfection with *mRFP-XK*. Scale bar = 5 μm. Images shown are representative of >100 cells examined for each transfection. (C) HEK293T cells were transfected with constructs expressing GFP-VPS13A^3027-3174^ (indicated by the blue box) or GFP-VPS13A^3027-3174^-I3148P (indicated by the blue box with an asterisk) with or without untagged *XK* and processed as in Figure 3A.

## Discussion

The paradigm developed in yeast is that Vps13 is recruited to specific membrane contact sites via interaction of the VAB domain with organelle-specific adaptor proteins (Bean et al., 2018; John Peter et al., 2017; Park et al., 2013). Point mutations in the VAB domain have been identified in patients with VPS13A, B, and D related diseases, indicating that this domain is critical for VPS13 function in human cells as well (Dobson-Stone et al., 2002; Gauthier et al., 2018; Kolehmainen et al., 2003). However it is becoming clear that Vps13 recruitment and function is more complex than this simple paradigm. For example, a recent report in yeast showed that the Arf1 protein binds to the PH domain of Vps13 to recruit Vps13 to the Golgi (Kolakowski et al., 2021).

This work has revealed that the PH domain of VPS13A plays a critical function in human cells by providing an interaction interface with XK. Unlike mutations in the VPS13A VAB domain that disrupt VPS13-XK co-localization without disrupting VPS13-XK complexes, mutants that truncate the PH domain in VPS13A change a conserved cytoplasmic loop in XK, or disrupt a predicted interaction surface between VPS13A and XK all eliminate both complex formation and co-localization. These results argue that the interaction between VPS13 and XK is direct and provide strong support for the idea that the phenotypic similarities between VPS13A disease and McLeod syndrome are due to disruption of this complex.

We propose that the presence of multiple partner binding sites reflects regulation of VPS13 protein localization via combinatorial recruitment. For example, there are at least three domains in VPS13A that are necessary for lipid droplet localization: (1) the VAB domain, (2) the PH domain and (3) the ATG-C domain (Kumar et al., 2018; Park and Neiman, 2020)(Figure 3B). VPS13A disease mutations in either the VAB or PH domain alone disrupt localization to lipid droplets, indicating that interaction between VPS13A and at least two other proteins is necessary for recruitment to lipid droplets.

The PH domain is required both for lipid droplet localization and for binding to XK. However, XK is unlikely to be directly responsible for recruitment to lipid droplets. XK itself is not localized at lipid droplets and, in fact, XK overexpression recruits VPS13A away from lipid droplets (Figure 2A; (Park et al., 2016)). Thus, analogous to the situation in yeast where multiple Vps13 adaptor proteins compete for binding to the VAB domain, our results suggest that XK competes with an unknown, lipid-droplet localized VPS13A partner for binding to the VPS13A PH domain.

The idea that multiple interactions regulate VPS13A association with different membranes is also supported by an Alphafold structural prediction of the full-length protein (Figure 6). Alphafold2 was used to generate a full-length model of the VPS13A-XK complex by dividing VPS13A into four slightly overlapping sections, predicting the structure of each section independently, and then aligning the overlapping regions of the structures (Figure 6). The XK protein was then docked onto VPS13A using the predicted structure of the interface shown in Figure 5A. The position of the transmembrane domains of XK indicate where the organelle membrane would be located and allow the orientation of the C-terminal end of VPS13A with respect to the membrane. Interestingly the VAB, ATG-C and PH domains in the C-terminal half of the protein are all in position to link the VPS13A to the membrane (Figure 6). The VAB domain and the PH domain likely do this by binding to membrane-associated proteins, e.g. adaptor proteins to the VAB domain and XK to the PH domain. Additionally, this structure predicts that the ATG-C domain consists of four amphipathic helices that expose a hydrophobic surface on the end of the protein (arrow in Figure 6), potentially constituting a direct membrane binding site. Consistent with this idea, the ATG-C domain in the yeast protein binds directly to lipids (De et al., 2017). Combinatorial binding through these three different motifs regulates the association of the C-terminal end of VPS13A with different membrane organelles, for example the mitochondrion, the lipid droplet, or the plasma membrane (see below) while the N-terminal end of VPS13A is anchored to the ER through binding to VAP family proteins (Kumar et al., 2018; Yeshaw et al., 2019).

**Figure 6.**
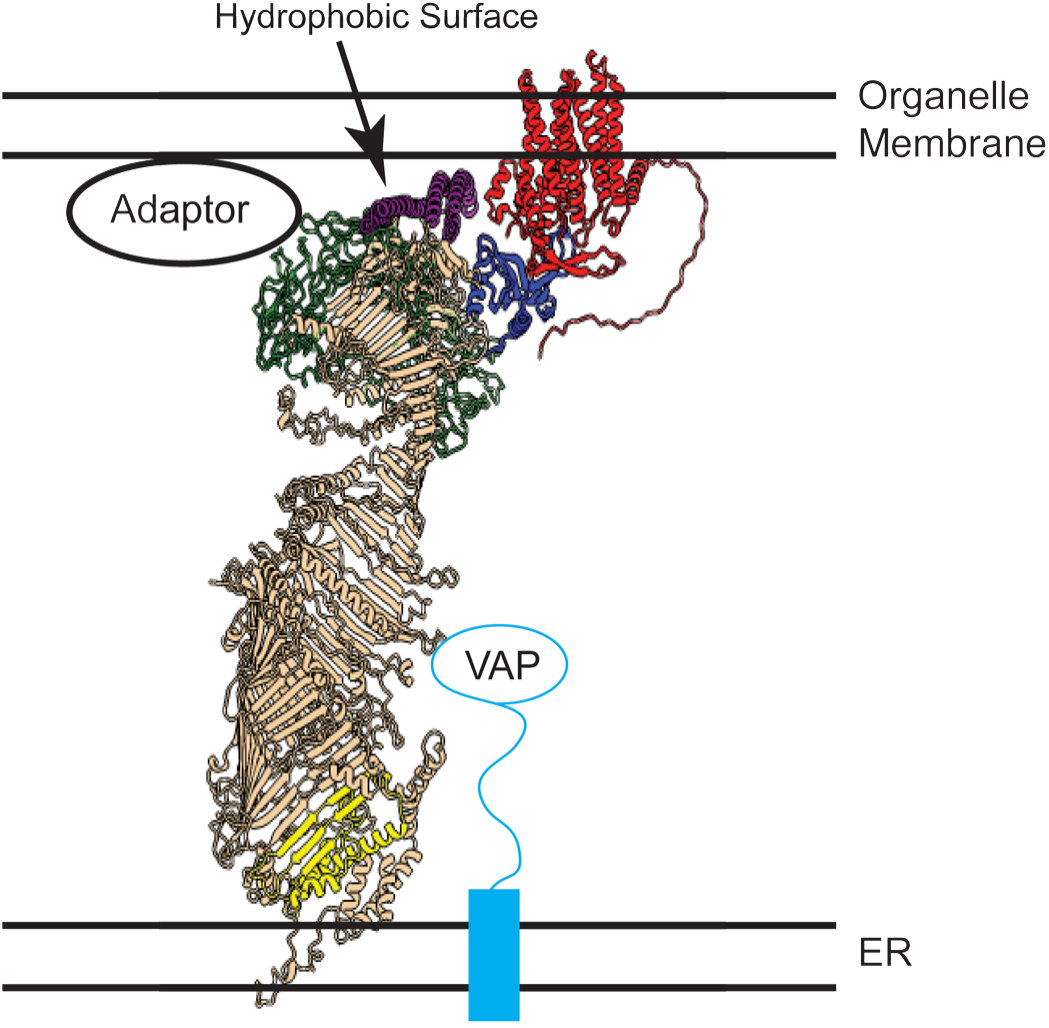
Model of VPS13A arrangement at a membrane contact site. A model structure of the VPS13A-XK complex based on assembly of Alphafold predictions is shown relative to the predicted locations of the ER membrane and a 2^nd^ organellar membrane (e.g. lipid droplet, mitochondria or plasma membrane). XK protein is in red and the different domains of VPS13A are highlighted: yellow for the N-Chorein domain, green for the Vps13 adaptor binding (VAB) domain, purple for the ATG2 C-terminal domain, and blue for the PH-like domain. Binding to the VAP protein anchors the C-terminal end of VPS13A to the ER (Yeshaw et al., 2019).

Our results are consistent with the idea that VPS13A disease and McLeod Syndrome both result from loss of an XK-VPS13A complex. An important question is where that complex is localized. We note that the colocalization of VPS13A and XK in ER subdomains, while a useful tool for examining association of the proteins, likely represents a consequence of overexpression of both proteins rather than the localization of the complex at native expression levels. In red blood cells, XK localizes to the plasma membrane as a component of the Kell antigen (Russo et al., 1999). While this work was in preparation, Guillen-Samander et al. (Guillen-Samander et al., 2022) reported mapping the interaction of VPS13A and XK to the PH domain and the second intracellular loop, respectively, consistent with the results presented here. Additionally, they demonstrated that VPS13A and XK can be co-localized at ER-plasma membrane junctions (Guillen-Samander et al., 2022). Thus, ER-plasma membrane junctions are a likely site of VPS13A-XK activity.

*XK* encodes a lipid scramblase, capable of flipping individual phospholipids between the leaflets of a membrane bilayer (Adlakha et al., 2022; Ryoden et al., 2022). XK is required for extracellular exposure of phosphatidylserine in response to extracellular ATP in immune cells and VPS13A is necessary for XK activity in this context (Ryoden et al., 2022). These results suggest that mediating exposure of phosphatidylserine on certain neurons may be the critical function of VPS13-XK, though XK scramblase activity is not specific to phosphatidylserine *in vitro* (Adlakha et al., 2022).

Physical interaction with lipid scramblases has emerged as a common property of VPS13-family and related lipid transport proteins (Adlakha et al., 2022; Ryoden et al., 2022). This linking of intermembrane and intrabilayer transport may be important for promoting transport through the VPS13 channel by maintaining a concentration gradient in lipids between the donor and acceptor leaflets. Knockdown of *VPS13A* in PC12 cells has been shown to lower phosphatidylinositol-4-phosphate levels at the plasma membrane (Park et al., 2015). Interestingly, maintenance of phosphatidylinositol-4-phosphate levels in the plasma membrane requires continual phosphatidylinositol synthesis in the ER, suggesting that continual delivery of phosphatidylinositol from the ER to the plasma membrane is required (Pemberton et al., 2020). Thus, an alternative possibility is that movement of lipids from the inner leaflet of the plasma membrane to the outer leaflet via XK helps maintain a directional flow of phosphatidylinositol from the ER to the plasma membrane via VPS13A. If true, then defects in phosphatidylinositol rather than phosphatidylserine transport could be the basis for the neurological effects of *VPS13A* and *XK* mutants. Further studies will be required to define the critical cellular role of VPS13A-XK that, when abolished, results in disease.

## Materials and Methods

### Cell culture and transfections

HEK293T cells were maintained at 37°C in a humidified atmosphere at 5% CO_2_ in Dullbecco’s Modified Eagle Medium (DMEM) supplemented with 10% Fetal Bovine Serum (FBS). Transfection for HEK293T cells was done using Lipofectamine 2000 (Thermo Fisher Scientific, cat. 11668-019) following the manufacturer’s instructions. Briefly, for microscopy, 2 μg plasmid DNA and 3μl Lipofectamine 2000 were separately added to 200 μl OptiMEM (Gibco, cat. 31985-062). After 5 minutes incubation at room temperature, the Lipofectamine 2000 mixed with OptiMEM was added to the plasmid DNA plus OptiMEM and this mixture was incubated at room temperature for 20 minutes. The plasmid/Lipofectamine 2000 mixture in OptiMEM medium was then added drop-wise onto HEK293T cells. For live cell microscopy, HEK293T cells were grown in gelatin coated 35-mm glass bottom culture dishes (Millipore, cat. ES-006-B). For immunoprecipitation and Western blot analysis, HEK293T cells were grown in gelatin-coated 10-cm culture dishes and 10 μg plasmid DNA and 20μl of Lipofectamine 2000 were used for transfection.

### Plasmid construction

Plasmids and primers used in this study are listed in Tables 1 and 2, respectively. Plasmid JS142-D3 expressing *mRFP-XK* was constructed by Gibson assembly using NEBuilder HiFi DNA Assembly (New England BioLabs, cat. E2611L) with an *mRFP* coding fragment amplified with oligos JSO620 and JSO628 and linearized pcDNA3.1(+)- N-*XK* as the template. To construct pJS160, which expresses *VPS13A^1-3143^^mCherry*, two fragments were amplified using the oligo pairs JSO731 / JSO732 and JSO733 / JSO734 with p*VPS13A*^mCherry as template. These primers amplify two overlapping halves of the template, creating a deletion of the 31 codons at the 3’ end of the *VPS13A* coding region. To construct plasmids expressing various *VPS13A* fragments fused to *GFP* each fragment was amplified with a pair of oligos by PCR; *VPS13A^2751-3174^* with JSO756 / JSO757, *VPS13A^3027-3174^* fragment with JSO775 / JSO757, *VPS13A^3027-3143^* with JSO775 / JSO781, and *VPS13A^3144-3474^*with JSO776 / JSO757 resulting in pJS164, pJS166, pJS171, and pJS167, respectively. The vector pEGFP-C2 was linearized by digestion with *Xho*I and *Xma*I and fragments were incorporated into the linearized vector by Gibson assembly. To create the point mutation I3148P within *VPS13A^3027-3174^*, the coding region for this *VPS13A* region was amplified in two overlapping fragments from pJS166 using the pairs of oligos JSO775/ JSO783 and JSO784 / JSO757. The mutation (‘ATA’ to ‘CCA’) I3148P was incorporated into the overlap between these two fragments. The two fragments then were introduced into the linearized vector pEGFP-C2 by Gibson assembly, leading to plasmid JS172.

**Table 1.**
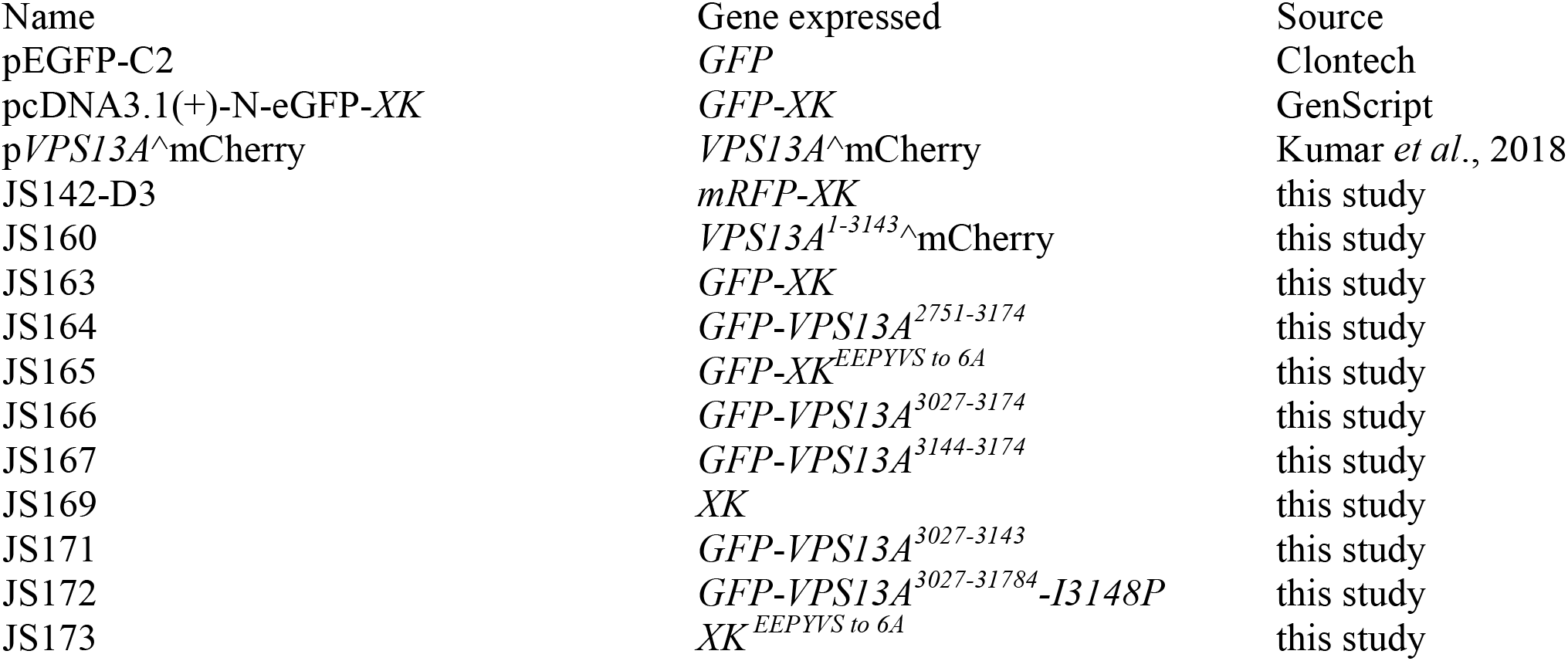
Plasmids used in this study

**Table 2.**
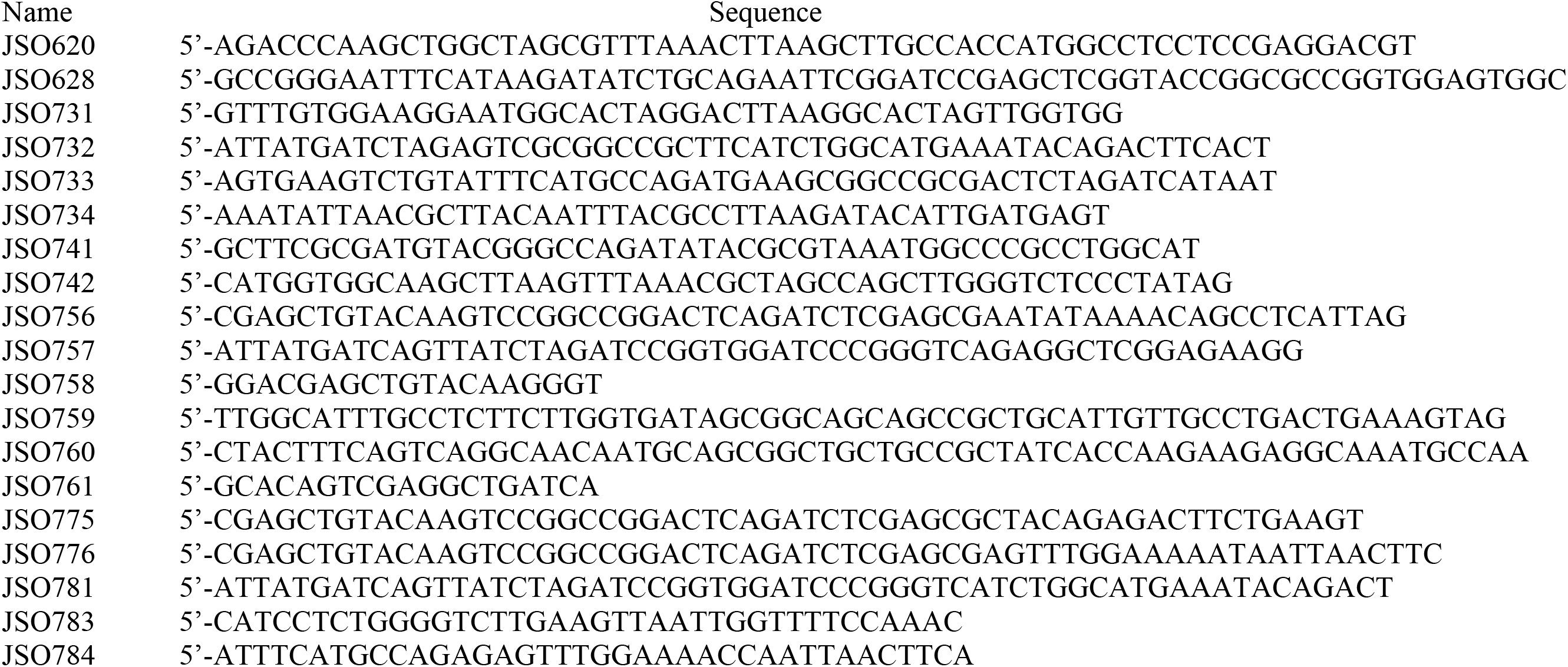
primers used in this study

To construct plasmid JS163, the full-length Cytomegalovirus (*CMV*) promoter was deleted from pcDNA3.1(+)-N-eGFP-*XK* by digestion with *Mlu*I and *Nhe*I and the vector backbone was purified using the Qiagen Gel purification kit (Qiagen, cat. 169021320). A smaller *CMV* promoter fragment (−125bp) was amplified from pcDNA3.1(+)-N-eGFP-*XK* with the oligo pair JSO741 / JSO742 and introduced into gel-purified pcDNA3.1(+)-N-eGFP-*XK* backbone by Gibson assembly. To generate plasmid JS165 which expresses *GFP-XK^6A^*, overlapping fragments were amplified from pcDNA3.1(+)-N-eGFP-*XK* with oligo pairs JSO758 / JSO759 and JSO760 / JSO76. These amplifications generate the 5’ and 3’ ends of the *XK* coding region with an overlap in the region of codons 90 to 110. In this overlap, the oligos alter the coding region for the amino acid sequence EEPYVS to AAAAAA (GAAGAGCCTTATGTCAGT to GCAGCGGCTGCTGCCGCT) pcDNA3(+)-N-eGFP-*XK* was digested with *EcoR*I and *Xba*I to remove the *XK* gene and the remaining backbone of the plasmid was assembled with the two PCR fragments by Gibson assembly. To remove the *GFP* tag from *GFP-XK* and *GFP-XK ^6A^*, pcDNA3.1(+)-N-eGFP-XK and pJS165 were first digested with *Hind*III and *EcoR*I to delete the *GFP* coding region. The 5’ and 3’ ends were filled in using DNA polymerase I Large (Klenow) fragment (New England Biolabs, cat. M0210S), and the blunt ends were ligated using T4 DNA ligase (New England Biolabs, cat. M0202S), resulting in pJS169 and pJS173.

Coding sequences from all constructed plasmids were confirmed by sequencing performed by the Stony Brook University DNA sequencing facility.

### Immunoprecipitation and Western blot analysis

Transfected cells were washed with ice-cold phosphate-buffered saline (PBS) (Life Technologies, cat. 14190-144), collected using a cell scraper into a 1.5 mL microcentrifuge tube, and stored at −80°C. GFP pull-downs were performed as described in (Park and Neiman, 2020) with minor changes. Cell pellets were resuspended in 300 μl lysis buffer (10 mM Tris/Cl pH7.5; 150 mM NaCl; 0.5 mM EDTA; 0.5% NP-40; 1 mM PMSF; and protease inhibitor cocktail (Thermo Scientific, cat. A32955; 1 tablet per 10 ml) and lysed by sonication with a microprobe (QSONICA Sonicators) at amplitude 40 for 15 sec, three times per sample. Cell lysates were centrifuged at 14,000 x g for 10 min at 4° C and supernatants were transferred into new 1.5-mL microcentrifuge tubes. 30 μl lysate per each supernatant was saved and mixed with 2 x SDS sample buffer (100 mM Tris, pH 6.8; 4% SDS; 20% Glycerol; 0.2 mg/ml Bromophenol blue; 0.72 M 2-Mercaptoethanol). To precipitate GFP-fusion proteins from the supernatants, 500 μl of washing buffer (10 mM Tris/Cl pH7.5; 150 mM NaCl; 0.5 mM EDTA) containing 1mM PMSF and protease inhibitor cocktail was added to the remaining supernatant to dilute the NP-40 down to 0.2%,, and then 25 μl of GFP-trap agarose beads (Chromotek, cat. GTA020), pre-washed twice with 500 μl washing buffer, were added. The supernatant/GFP-trap agarose bead mixtures were incubated for 1.5 hrs at 4° C on a rotator. Agarose beads were pelleted by centrifugation at 2,500 x g for 2 min at 4°C. Beads were washed twice with 500 μl washing buffer. Finally, 2 x SDS sample buffer was added to the beads and samples were heated at 50° C for 10min.

Protein samples were fractionated on 7.5% or 10% SDS-polyacrylamide gels (Bio-Rad, cat. 1610171 or 1610173). For the transfer of VPS13A full-length protein tagged with or without mCherry, transfer of the 7.5% gel onto PVDF membrane was done overnight at 4° C as described in Park *et al*. (2013). Primary antibodies used for immunoblot analysis were α-VPS13A (Sigma, cat. HPA021652) at 1:700 dilution, α-mCherry (Rockland, cat. 600-401-P16) at 1:1000 dilution, α-GFP (cat. 632381; Clontech) at 1:1000 dilution, α-XK (Sigma, cat. HPA019036) at 1:2500 dilution, and α-GAPDH (Proteintech, cat. 60004) at 1:5000 dilution. Secondary antibodies were ECL anti-mouse IgG (GE Healthcare, cat. NA931V) at 1:5000 dilution and ECL anti-rabbit IgG (GE Healthcare, cat. NA934V) at 1:5000 dilution.

### Preparation of and Western blotting of human blood samples

2,5 ml aliquot of EDTA/Citrate blood was defrosted, filled up to 12 ml with RBC wash buffer (5 mM Na_2_HPO_4_, 154 mM NaCl), mixed carefully and centrifuged for 10 min at 4000 rpm. Supernatant was discarded. 1.5-2 ml pellet was left at the bottom. Washing steps were repeated for pellet. Supernatant was removed and discarded. Pellet was distributed to two 1,5 ml standard lab reaction tubes. 500 µl RBC lysis buffer (5 mM Na_2_HPO_4_) was added to each reaction tube. Samples were centrifuged for 5 min at 13000 rpm and then supernatant was carefully removed and discarded. Samples was washed with RBC lysis buffer for at least 2 times. After that, samples were distributed in 5 µl aliquots and stored at −20 °C.

For immunoblots of these samples, 15 µl 2× Lämmli buffer was added to 5 µl aliquots of blood samples. Samples were incubated at 70 °C for 10 minutes. A total volume of 10 µl per lane was loaded to NuPAGE^TM^ 3-8% Tris-Acetate gel (Invitrogen, Cat. EA0375BOX). After electrophoresis, probes were transferred to PVDF membranes. Membranes were blocked for 1 h in PBST with 0.2% I-Block (Invitrogen, T2015). The membranes were incubated overnight with primary antibody in block solution at 4 °C. After washing 4 × 15 min with PBST, the membranes were incubated with secondary antibodies for 1 h in PBST with 0.2% I-Block. Development of the membrane after 4 × 15 min PBST washes was performed with ECL plus. The α-VPS13A antibody (Sigma, cat. HPA021662) was used at 1:10,000 dilution and detected with HRP conjugated α-rabbit antibodies (Promega, W4011) used at 1:10,000 dilution. ß-actin was detected using primary antibodies (Sigma, cat. A5316) at 1:50,000 dilution and HRP conjugated α-mouse antibodies (Promega, cat W4021) at 1:5,000 dilution.

### Immunostaining and microscopy

To visualize lipid droplets in live cells, 70∼80% confluent cells in gelatin-coated 35 mm glass bottom culture dishes were washed once with Hank’s balanced salt solution (HBSS) (cat. 14025092; Life Technologies) buffer and incubated with 1 mL HBSS buffer containing monodansylpentane (MDH) (cat. SM1000a; ABCEPTA) at 5 μM for 15 min in a 37° C humidified incubator. To determine fluorescent protein localizations in live cells, cells at 24 hours after transfection were washed once and incubated with HBSS buffer. Live cells within HBSS buffer were observed using a wide-field Zeiss Observer.A1 microscope with an attached Orca II ERG camera (Hamamatsu) or a Zeiss LSM 510 META NLO Two-Photon Laser Scanning Confocal microscope coupled with a Coherent Chameleon laser system. ZEN 2012 Blue edition software was used to acquire and process images from the wide-field microscope. Zeiss LSM version 4.0 software and ZEN Black edition software were used to acquire and process images from the confocal microscope, respectively.

## Acknowledgements

The authors wish to acknowledge the work of Professor Adrian Danek, Dr. Benedikt Bader (both Department of Neurology at Ludwig-Maximilians-Universität Munich, Germany) and Mrs. G. Kwiatkowski (Department of Neuropathology, Ludwig-Maximilians-Universität Munich, Germany) in establishing the repository for the patient samples used is this study, as well as the support of the Advocacy for Neuroacanthocytosis Patients, the ERA-net E-Rare consortium EMINA (European Multidisciplinary Initiative on Neuroacanthocytosis; BMBF01GM1003) and all the collaborating clinicians who sent samples for analysis. We are grateful to Bettina Schmid and Adrian Danek for comments on the manuscript. We also thank the Stony Brook Central Microscopy Facility for access to the confocal microscope and members of the Neiman Lab for helpful discussion.

